# Dysregulation of lung epithelial cell homeostasis and immunity contributes to Middle East Respiratory Syndrome coronavirus disease severity

**DOI:** 10.1101/2024.10.03.616483

**Authors:** Amy C. Sims, Alexandra Schäfer, Kenichi Okuda, Sarah R. Leist, Jacob F. Kocher, Adam S. Cockrell, Kara L. Jensen, Jennifer E. Kyle, Kristin E. Burnum-Johnson, Kelly G. Stratton, Natalie C. Lamar, Carrie D. Niccora, Karl K. Weitz, Richard D. Smith, Thomas O. Metz, Katrina M. Waters, Richard C. Boucher, Stephanie A. Montgomery, Ralph S. Baric, Timothy P. Sheahan

**Affiliations:** Department of Epidemiology, University of North Carolina at Chapel Hill, Chapel Hill, NC USA; Marsico Lung Institute, University of North Carolina at Chapel Hill, Chapel Hill, NC, USA; Biological Sciences Division, Pacific Northwest National Laboratories, Richland, WA USA; AI & Data Analytics Division, Pacific Northwest National Laboratories, Richland, WA USA; Department of Pathology & Laboratory Medicine, University of North Carolina, Chapel Hill, NC; Department of Microbiology and Immunology, University of North Carolina at Chapel Hill, Chapel Hill, NC USA; Nuclear, Chemical and Biological Technologies Division, Pacific Northwest National Laboratory, Richland, WA USA

## Abstract

Coronaviruses (CoV) emerge suddenly from animal reservoirs to cause novel diseases in new hosts. Discovered in 2012, Middle East respiratory syndrome coronavirus (MERS-CoV) is endemic in camels in the Middle East and is continually causing local outbreaks and epidemics. While all three newly emerging human CoV from past 20 years (SARS-CoV, SARS-CoV-2, MERS-CoV) cause respiratory disease, each CoV has unique host interactions that drive differential pathogeneses. To better understand the virus and host interactions driving lethal MERS-CoV infection, we performed a longitudinal multi-omics analysis of sublethal and lethal MERS-CoV infection in mice. Significant differences were observed in body weight loss, virus titers and acute lung injury among lethal and sub-lethal virus doses. Virus induced apoptosis of type I and II alveolar epithelial cells suggest that loss or dysregulation of these key cell populations was a major driver of severe disease. Omics analysis suggested differential pathogenesis was multi-factorial with clear differences among innate and adaptive immune pathways as well as those that regulate lung epithelial homeostasis. Infection of mice lacking functional T and B-cells showed that adaptive immunity was important in controlling viral replication but also increased pathogenesis. In summary, we provide a high-resolution host response atlas for MERS-CoV infection and disease severity. Multi-omics studies of viral pathogenesis offer a unique opportunity to not only better understand the molecular mechanisms of disease but also to identify genes and pathways that can be exploited for therapeutic intervention all of which is important for our future pandemic preparedness.

**Importance:** Emerging coronaviruses like SARS-CoV, SARS-CoV-2 and MERS-CoV cause a range of disease outcomes in humans from asymptomatic, moderate and severe respiratory disease which can progress to death but the factors causing these disparate outcomes remain unclear. Understanding host responses to mild and life-threatening infection provides insight into virus-host networks within and across organ systems that contribute to disease outcomes. We used multi-omics approaches to comprehensively define the host response to moderate and severe MERS-CoV infection. Severe respiratory disease was associated with dysregulation of the immune response. Key lung epithelial cell populations that are essential for lung function get infected and die. Mice lacking key immune cell populations experienced greater virus replication but decreased disease severity implicating the immune system in both protective and pathogenic roles in the response to MERS-CoV. These data could be utilized to design new therapeutic strategies targeting specific pathways that contribute to severe disease.

## Introduction

The epidemic and pandemic potential of the family *Coronaviridae* (CoV) is evidenced by the emergence of severe acute respiratory syndrome coronavirus (SARS-CoV) in 2002, Middle East respiratory syndrome CoV (MERS-CoV) in 2012 and SARS-CoV-2, the etiologic agent of the COVID-19 pandemic, in 2019 (1–3). These emerging CoVs primarily cause respiratory disease that range from subclinical asymptomatic disease to end stage acute lung injury (ALI), acute respiratory distress syndrome (ARDS) and death. To date, MERS-CoV has caused 2613 lab-confirmed cases in 27 countries with a case fatality rate of 39% (4). Most cases have occurred in the Kingdom of Saudi Arabia. Serological surveys suggest mild disease presentations are common and that case numbers are underestimated (5). MERS-CoV is also endemic in dromedary camels in the Middle East and Northeast Africa with serologic evidence of infection going back three decades (6). While most human MERS-CoV cases have been localized to the Middle East, air travel of an infected person caused the 2015 outbreak of 186 cases with 38 deaths in South Korea (7, 8). The MERS-CoV reproductive rate (R_0_), or the number of secondary transmissions associated a primary infection, is less than one (0.9) indicating that most primary infections fail to cause secondary infections (9). In MERS-CoV and SARS-CoV 2003, superspreaders were rare but also contributed significant to outbreak expansions (10). In contrast, the average R_0_ for SARS-CoV-2 Omicron is 9.5, which is indicative of efficient to explosive person to person transmission (11). The cellular entry receptor for MERS-CoV, dipeptidyl peptidase 4 (DPP4), is expressed in multiple tissues and cell types including lung epithelial cells and immune cells (12, 13). In humans, DPP4 expression in the lung is mostly limited to the lower airways yet in camels receptor expression is pervasive throughout the upper and lower airway suggesting that the lack of upper airway expression in humans limits efficient transmission (12). While most MERS-like CoV (i.e. *Merbecoviruses*) utilize DPP4 for entry, angiotensin converting enzyme 2 (ACE2), the receptor for most SARS-like CoVs (i.e. *Sarbecoviruses*), has recently been identified as the entry receptor for some enzootic *Merbecoviruses* (14). As evidenced by SARS-CoV-2 and human CoV 229E, CoVs can rapidly evolve increased human transmissibility, but to date, MERS-CoV has undergone relatively little genetic change and remains inefficiently transmitted from person to person (15).

Despite considerable study, the host determinants that regulate MERS-CoV disease severity are not completely understood. The first human autopsy case from a MERS-CoV infected patient, published three years after the virus’ discovery, confirmed infection was associated with pneumonia, diffuse alveolar damage and infiltrating inflammatory cells (16, 17). MERS-CoV primarily targets lung alveolar type I (AT1) and II (AT2) pneumocytes for infection although most human and non-human primate studies have identified infected cells based only on morphology and the host responses in these cells and the consequences of their infection remains unclear (16–19). Using primary cells from human lungs, efficient MERS-CoV replication has been documented in non-ciliated airway epithelial cells, AT2 cells, lung fibroblasts and endothelial cells with cell type specific host expression patterns (20, 21). Since MERS-CoV cannot utilize the murine ortholog of DPP4 (mDPP4) for entry, we created a mouse model by introducing “humanizing” mutations A288L and T330R in mDPP4 in C57BL/6 mice (C57BL/6 hDPP4) (22). Here, we used a multi-omics approach to comprehensively define the host response to sub-lethal and lethal MERS-CoV infection in C57BL/6 hDPP4 transgenic mice (22). These data can be utilized to better understand the similarities and differences in the host response patterns across highly pathogenic emerging CoV and can be harnessed to develop host targeting antivirals that impact pathways commonly leveraged by CoV for replication and disease.

## Results

### MERS-CoV infection elicited acute lung injury is virus dose dependent

To broadly define the virologic, pathologic, transcriptomic, proteomic and lipidomic signatures associated with sublethal and lethal MERS-CoV infection, we intranasally inoculated age and sex matched C57BL/6 hDPP4 mice with PBS (Mock), 5×10^4^ plaque forming units (PFU) (Low Dose, sublethal) or 5×10^6^ PFU (High Dose, lethal) mouse adapted MERS-CoV MA1 strain (22). After infection, body weights were measured daily and tissue samples were harvested on 2, 4 and 7 days post infection (dpi) for virology, histology, transcriptomics, proteomics and lipidomics analysis (Fig. 1A) (22). Weight loss is a marker of emerging CoV pathogenesis in mice and as expected, mock infected mice did not lose weight during our study (Fig. 1B). In contrast, MERS-CoV infected mice exhibited virus dose-dependent body weight loss over time (Fig. 1B). Those receiving High Dose MERS-CoV lost significantly more weight than those infected with Low Dose (Fig. 1B). Similarly, after 7 days, 100% of Low Dose infected mice had survived while 90% of the High Dose group had met humane endpoints for euthanasia (Fig. 1C). Both dose groups had similarly high virus titers on 2dpi and 4dpi, but unlike the Low Dose group which had largely cleared virus by 7dpi (Fig. 1D. Low Dose mean 7dpi titer = 1.25×10^2^ PFU/lobe), the High Dose group failed to clear virus and high viral loads remained in the lung (Fig. 1D. High Dose mean 7dpi titer = 1.5×10^7^ PFU/lobe). Trends in infectious viral titers were mirrored in MERS-CoV antigen labeling of lung tissue sections (Fig. 1S). Lung discoloration, a gross pathological feature of CoV pathogenesis (23–26), was significantly increased in High Dose infected animals at later times post infection as compared with the Low Dose group (Fig. 1E). Together, these data demonstrate mouse adapted MERS-CoV disease severity is guided by virus dose and that our experimental conditions for our multi-omics study provided starkly different outcomes.

**Figure 1:**
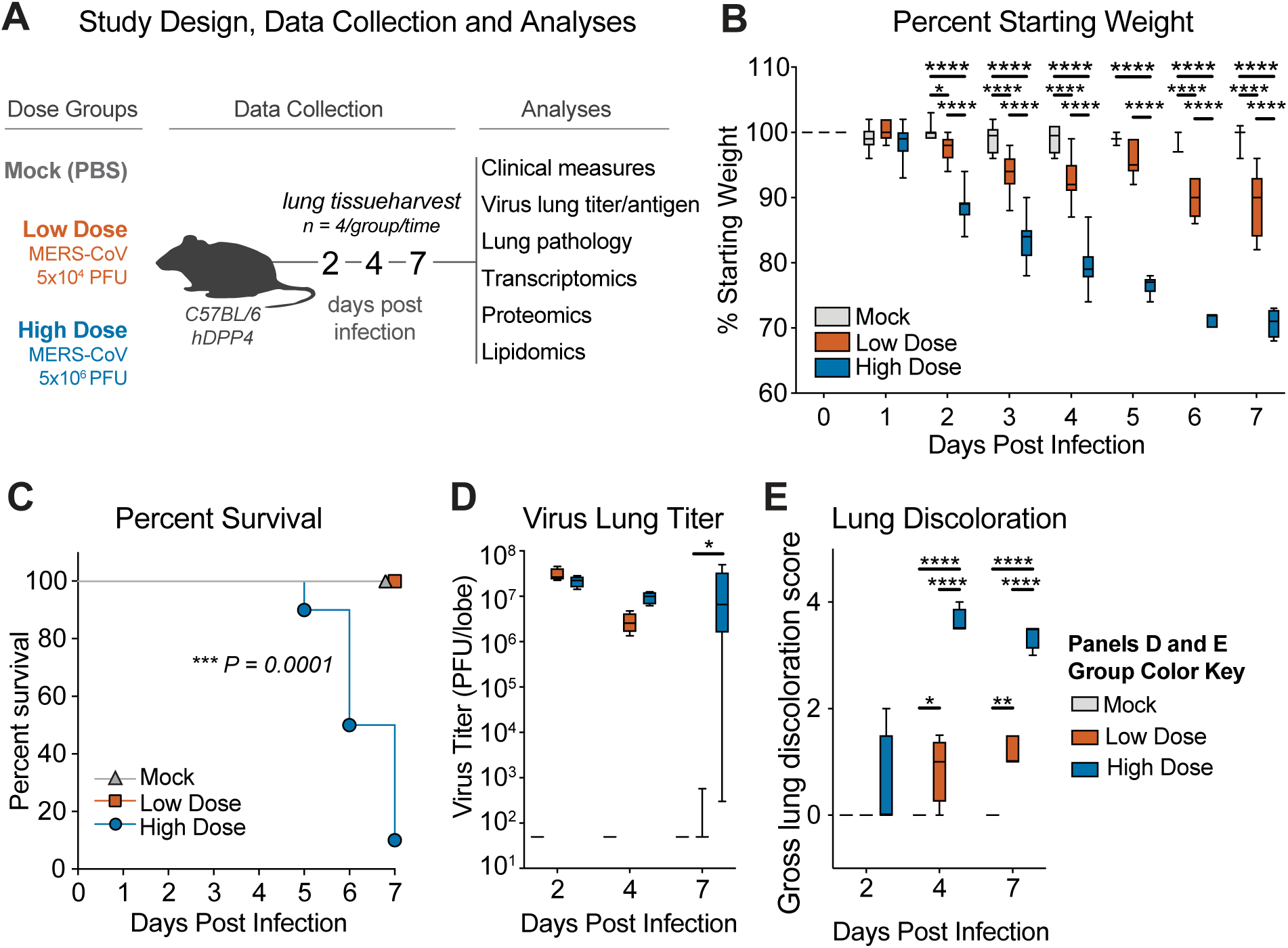
The host response and MERS-CoV disease severity is dose dependent. **(A)** Study design. 19-23 week old female C57BL/6 hDPP4 mice were infected with PBS (“mock”, N = 14), 5×10^4^ PFU mouse adapted MERS ma1 (“low dose”, N = 14) or 5×10^6^ PFU MERS ma1 (“high dose”, N = 16). On 2, 4, and 7dpi, lung tissue from 4 mice per group was harvested for virological, pathological, and multi-omics measures. **(B)** Percent starting weight. The boxes encompass the 25th to 75th percentile, the line is at the median, while the whiskers represent the range. Asterisks denote statistical significance as determined by Two-way ANOVA with a Tukey’s multiple comparison test. * = 0.01, **** = < 0.0001. **(C)** Percent Survival and significance as determined by Mantel-Cox test (***P = 0.0001). **(D)** Virus lung titer per lung lobe by plaque assay. Asterisks denote statistical significance by Two-way ANOVA with a Tukey’s multiple comparison test. * = 0.01. **(E)** Lung hemorrhage scored on a scale of 0–4, where 0 is a normal pink healthy lung and 4 is a completely dark red lung. Asterisks denote statistical significance as determined by Two-way ANOVA with a Tukey’s multiple comparison test. * = 0.01, ** = 0.001, **** < 0.0001.

MERS-CoV can cause diffuse alveolar damage (DAD), the pathological hallmark of acute lung injury (ALI), in humans and in mice (1, 25). We blindly evaluated hematoxylin and eosin-stained lung tissue sections for the histologic features of DAD/ALI. Lung tissues from mock infected mice appeared normal with thin alveolar septae suitable for efficient gas exchange and terminal airspaces free of debris and inflammatory cells (Fig. 2A). Although some features of ALI were noted in tissue from the Low Dose group (Fig. 2B), these features were far more prominent, especially at later times post infection, in the High Dose group including septal wall thickening, scattered dying and degenerating cells, proteinaceous debris and neutrophils in the air spaces (Fig. 2C). Interestingly, hyaline membranes, a proteinaceous alveolar lining formed in response to epithelial damage and vascular leakage, were noted in the Low Dose group (Fig. 2B) but were far more extensive in High Dose animals (Fig. 2C). We then quantitated the features of ALI/DAD using two complementary histologic scoring tools we recently validated in multiple mouse models of emerging CoV pathogenesis (Fig. 2D) (25–27). The American Thoracic Society Lung Injury scores were significantly increased and approached maximal levels in the High Dose MERS infected mice as compared to Low Dose MERS-CoV infected mice (Fig. 2D) (28). Similar trends were observed using a second scoring tool focused specifically on DAD phenotypes (Fig. 2D). Thus, severe MERS-CoV infection was associated with histologic features of DAD/ALI like those observed in human MERS-CoV cases (17).

**Figure 2:**
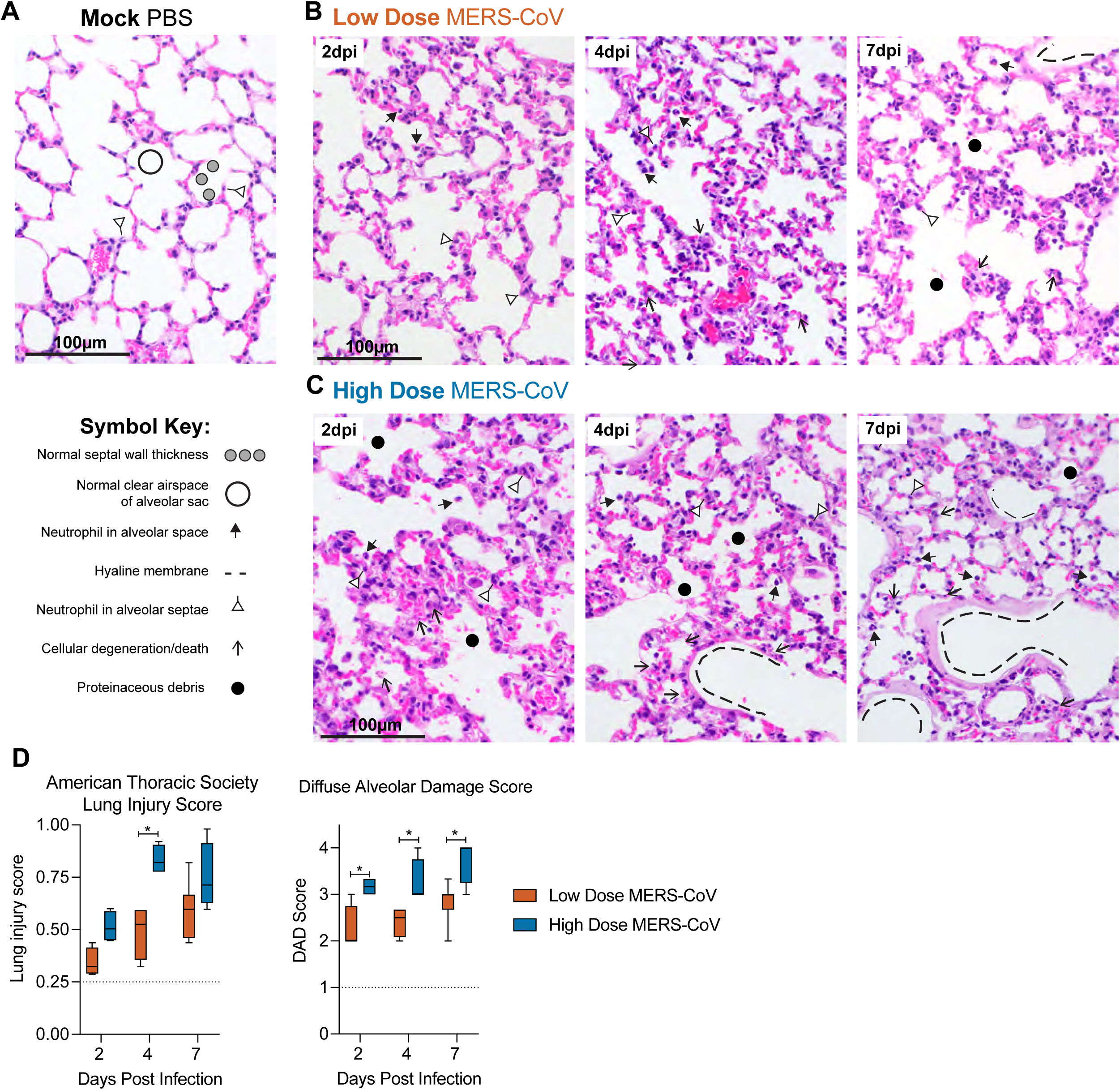
Severe MERS-CoV pathogenesis is associated with increased acute lung injury. For mock **(A)**, Low **(B)** and High dose **(C)** MERS-CoV infected groups, the histological features of acute lung injury were evaluated and blindly scored for: neutrophils in the alveolar and interstitial space, hyaline membranes, proteinaceous debris filling the air spaces, and alveolar septal thickening. Example features are noted according to the symbol key. **(D)** Features of acute lung injury were quantitated using an American Thoracic Society lung injury scoring system (Left) and a diffuse alveolar damage system (Right). Scores were blindly assessed in three random high power (×60) fields of diseased lung tissue sections. Mock N = 3; Low Dose 2 and 4dpi (N = 4/time), 7dpi (N = 7); High Dose N = 4/time for all time points. Th dotted line indicates the average scores observed in mock infected animals. Asterisks denote statistical significance as determined by Two-way ANOVA with Sidak’s multiple comparison test.

### Type I and type II epithelial cells infected with MERS-CoV undergo apoptosis

The cells MERS-CoV targets in humans and the impact of infection on cellular function is complicated by there being very few pathology reports from MERS-CoV patients (16, 17). MERS-CoV antigen has been found in bronchiolar, AT1 and AT2 epithelial cells in non-human primates and after infection of human lung ex-plant cultures (29, 30). AT1 cells line most of the alveolar air surface and form the air-blood barrier where gas exchange occurs (31). AT2 cells are a major producer of surfactant, a complex of lipids and protein which helps maintain surface tension and inflation of the lung (32). To determine if infection and cell death of key epithelial cell populations could be driving DAD, we utilized in situ hybridization and antigen staining to identify cell types targeted by MERS-CoV. We found MERS-CoV targeted a variety of epithelial subtypes in the distal airways, including both AT1 (*Ager* RNA+) and AT2 (*Sftpc* RNA+) cells (Fig. 3A), as well as *Scgb1a1* RNA positive secretory club cells in the conducting airways (Fig. 3B). To determine if infection resulted in apoptosis, we performed immunostaining for cell type and viral specific markers as well as cleaved caspase 3, a marker of apoptosis. MERS-CoV infection of AT1 (Fig. 3C) and AT2 (Fig. 3D) cells resulted in the induction of apoptosis as evidenced by the co-staining of cleaved caspase 3. Moreover, in situ hybridization for *Sftpc* transcript showed a dramatic loss of AT2 cells in High Dose MERS-CoV infected animals which were not replenished by the end of our study (Fig. 2S).

**Figure 3:**
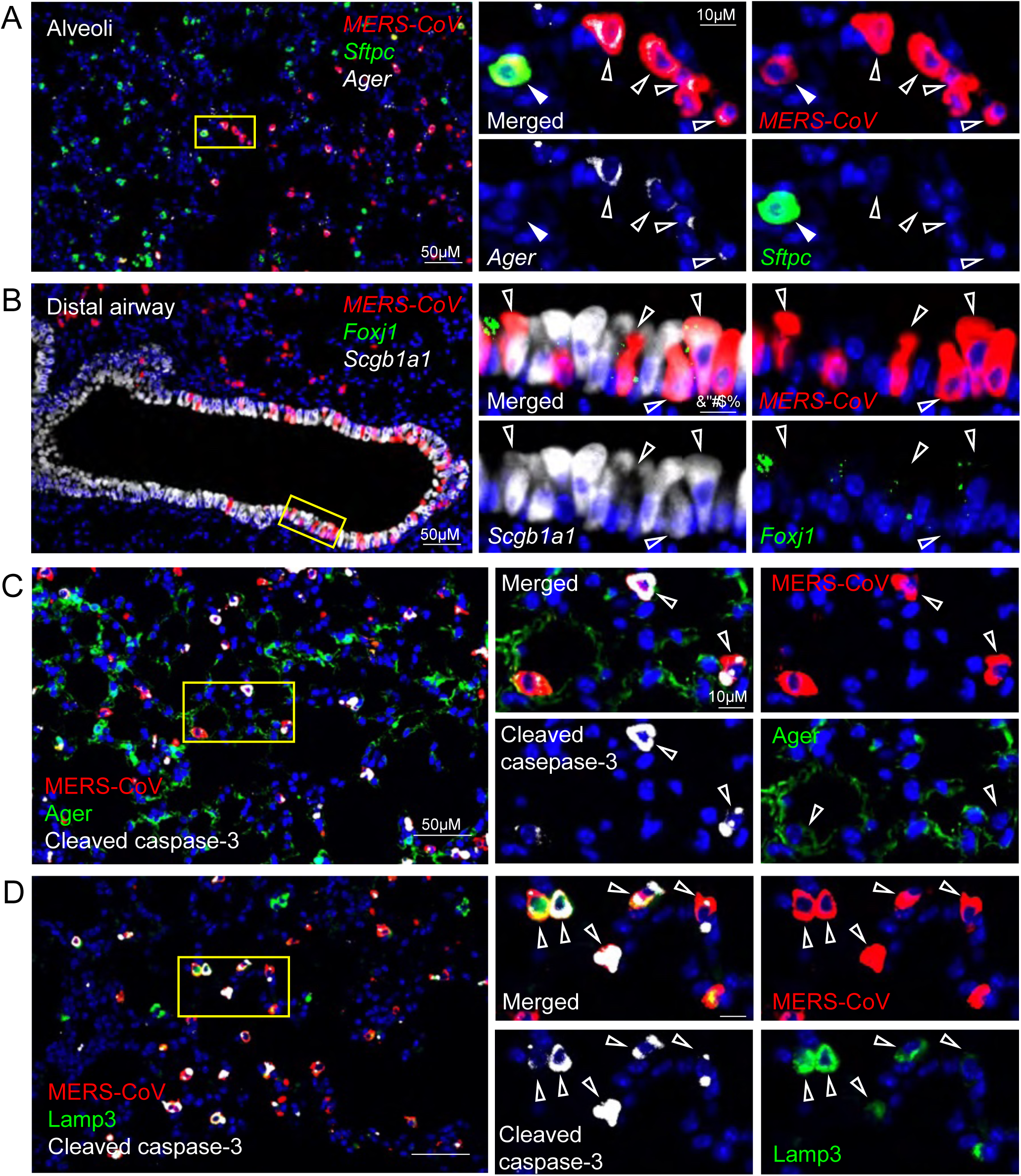
MERS-CoV infection is associated with apoptosis of type 1 and type 2 pneumocytes. MERS-CoV cellular tropism determine by in situ hybridization and immunohistochemistry. **(A)** RNAscope in situ hybridization for MERS-CoV RNA, *Ager* (advanced glycosylation end-product specific receptor mRNA, alveolar type I marker) and *Sftpc* (surfactant protein C mRNA, alveolar type II cell marker). **(B)** RNAscope in situ hybridization for MERS-CoV RNA, *Foxj1* (forkhead box J1, ciliated epithelial cells), and *Scgb1a1* (secretoglobin family 1A member 1, club cell marker). **(C)** Immunohistochemistry for MERS-CoV antigen, type I cell marker, AGER and apoptosis marker, cleaved caspase-3. **(D)** Immunohistochemistry for MERS-CoV antigen, type II cell marker, LAMP3 and apoptosis marker, cleaved caspase-3.

In agreement with data above, the impact of epithelial cell infection and apoptosis on the lung transcriptome, proteome and lipidome was notable. We filtered our transcriptomic and proteomic data using known lung epithelial cell signatures from single cell data sets (33). The expression of multiple genes related to AT1 (e.g. *AGER*) and AT2 (e.g. *LAMP3*) cells were significantly reduced on 2, 4 and 7 dpi in animals infected with High Dose MERS-CoV (Fig. S3A). Proteins associated both AT1 and AT2 cells were significantly downregulated in animals with severe MERS-CoV infection including key surfactant proteins synthesized by AT2 cells (e.g. SFTPA, SFTPB, SFTPC). Since the majority of surfactant is comprised of lipids, we next sought to determine the effect of severe MERS-CoV disease on the lung lipidome. In general, shorter chain phosphatidylcholine (PC e.g. PC(14:0/16:1); PC(16:0/16:1)) and phosphatidylglycerol (PG e.g. PG(16:0/18:1)) lipids, the main classes of surfactant lipids, were diminished with sub-lethal and severe MERS-CoV disease (Fig. S3C and D). In addition, shorter chained triacylglyceride (TG) lipids, typically found in lung resident mesenchymal cells which serve as a fatty acid reservoir for AT2 cells for surfactant lipid production (32), were similarly reduced in mice with severe MERS-CoV disease (Fig. S3C and D). These data coupled with the histologic data suggest that severe MERS-CoV infection results in a dysregulation of epithelial cell transcriptomic, proteomic and lipidomic programs essential for maintaining lung homeostasis. Together, we show that MERS-CoV infection and apoptosis of AT1 and AT2 cells is associated with the loss of epithelial barrier integrity, leakage of plasma protein components into the airway and ALI which complicates survival and recovery.

### Severe MERS-CoV disease is associated immune pathway dysregulation

To better understand the innate and adaptive immune responses associated with MERS-CoV induced ALI, we compared the lung transcriptomic and proteomic profiles from mock, Low Dose and High Dose MERS-CoV infected mice. Ingenuity IPA pathway analysis of transcriptomic data revealed multiple canonical pathways related to innate immune and adaptive immune responses were predicted to be differentially regulated among our two dose groups (Fig. 4). Pathways associated the innate antiviral response (e.g. acute phase response, recognition of viruses by pattern recognition receptors (PRR), etc.) were predicted to have increased function following Low dose MERS-CoV but decreased function with High dose MERS-CoV (Fig. 4A). Upon further inspection of pathways related to Interferon Signaling (Fig. 4SA), Recognition of Viruses by PRR (Fig. 4SB) and Interferon Stimulated Genes (ISGs, Fig. 4SB) and Proteins (4SC), the patterns of gene and protein expression were remarkably concordant among virus doses. For example, expression patterns for *Ifit3*, *Irf1*, *Irf9*, *Isg15*, and *Stat2* (Fig. 4SA) were very similar among virus doses much of which was similarly reflected in the proteome. Interestingly, in “Recognition of Viruses by PRR” pathway (Fig. 4SB), both virus doses induced expression of classic innate antiviral sensors (*Ddx58, Oas, Eif2aK2, Tlr3*), adaptor proteins (*MyD88*) and transcription factors (*Irf7*) but only severe disease was associated with the downregulation of several genes associated with the inflammatory response (*Fgrf4, Irs-1, C5, Prkcz*) (34–37) (Fig. 4SB). While the pattern of ISG gene and protein expression was for the most part comparable among high and low dose MERS-CoV infection, several ISGs with roles in the adaptive immune response were uniquely downregulated with high dose MERS-CoV infection. For example, *Cd74* (e.g. critical for antigen presentation on MHCII), *Nrn1* (e.g. enhances suppressive activity of regulatory T cells), and *Lppr* (e.g. aids in antigen specific adaptive immune responses) were uniquely downregulated in severe MERS-CoV disease (38–40). Altogether, these data demonstrate that differences in disease severity were not associated with major differences in the innate immune response.

**Figure 4:**
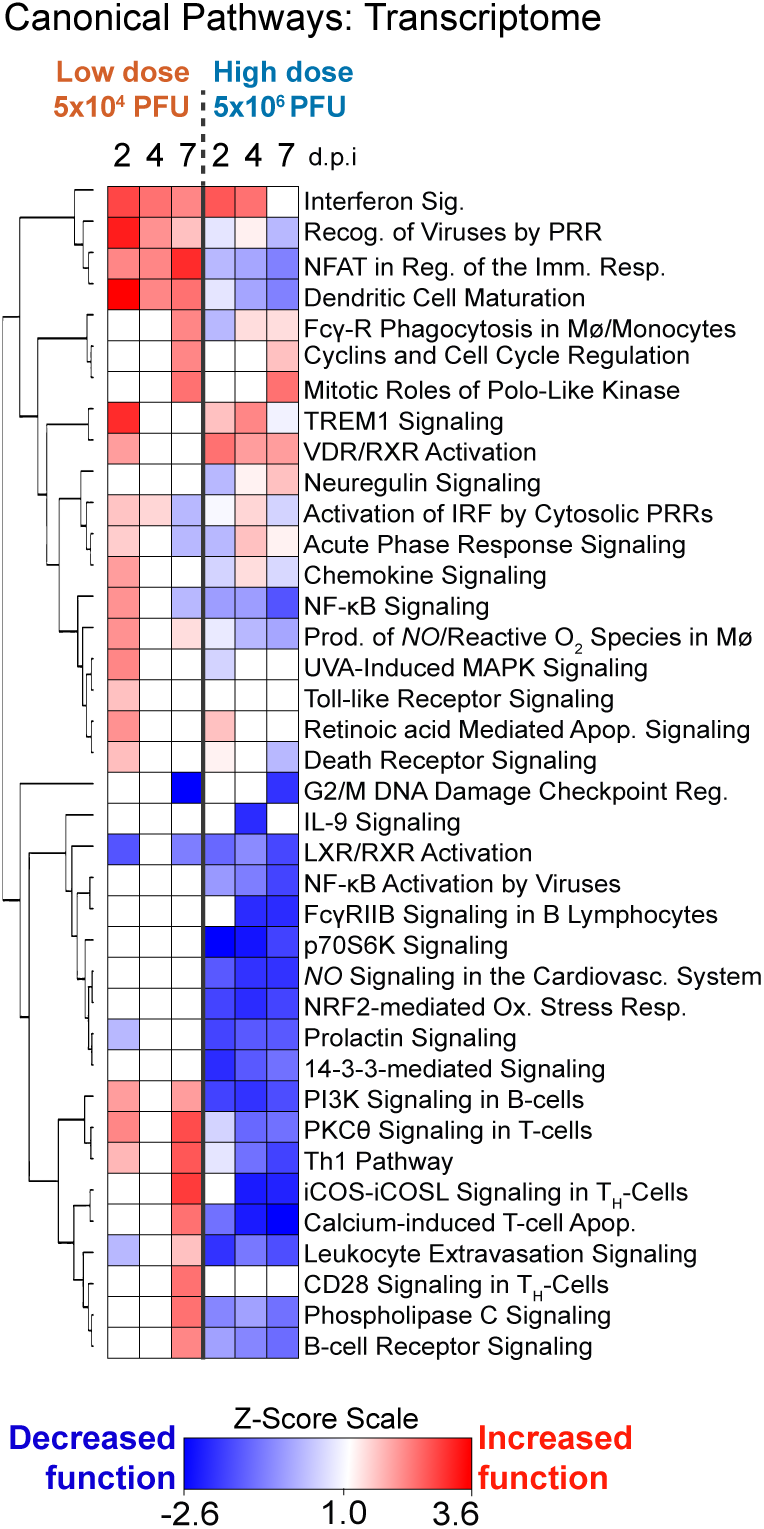
MERS-CoV disease severity is associated with differential regulation of innate and adaptive immune pathways. The Ingenuity IPA “canonical pathways” modulated in the significantly regulated host transcriptome by MERS-CoV over time. The heat map shows the predicted functional effect on canonical pathways (Z-scores > 1.5 = significant pathway activation and < −1.5 = significant pathway suppression).

Unlike the more subtle differences observed among Low Dose and High Dose MERS-CoV groups in the innate response, stark differences were noted in the predicted function of adaptive immune related pathways such as Th1 Pathway, NFAT Regulation of the Immune Response, Leukocyte Extravasation Signaling and p70S6K Signaling (Fig. 4). In Low Dose infected animals, unique expression of key adaptive immune genes (Fig. 5SA, *Cd3, Cd8, Stat4, Icos, Nfatc2*, etc.) was observed while unique suppression multiple major histocompatibility (MHC) genes was noted with High Dose MERS-CoV. Relatedly, downregulation of both gene and protein expression of multiple members of the NFAT Regulation of the Immune Response Pathway (*Cd79A/B*, B-cell receptor; *H2-Aa, H2-Ab1, H2-Eb1*, etc.) was associated with severe MERS-CoV pathogenesis (Fig. 5SB). Similarly, a downregulation of genes and proteins related to Leukocyte Extravasation Signaling (Fig. 5SC) and p70 S6K Signaling (Fig. 5SD) pathways, which are important for T or B cell related responses, was noted in the lung tissues of animals infected with High Dose MERS-CoV. Overall, the multi-omics data demonstrated a dysregulation in the adaptive immune response with severe MERS-CoV disease. We confirmed these molecular observations with multi-antigen immunostaining (Fig. 5). Lung tissue sections from mock, Low Dose and High Dose MERS-CoV infected mice were co-stained with MERS-CoV antigen and markers of key immune cell populations (i.e. CD68 monocyte/macrophage marker; CD4 or CD8, T cell subpopulation markers; B220, B cell marker). In Low Dose infected animals, there was an inverse correlation among the abundance of MERS-CoV antigen staining and CD68, CD4, CD8 and B220 positive cells (Fig. 5) as there was a general increase in immune marker staining over time. In contrast, MERS-CoV antigen remained abundant at all times assessed in lung tissue sections from High Dose MERS-CoV infected animals. Remarkably, aside from slight increases in CD4 positive cells on 7dpi, relatively little difference among mock and infected animals was observed in immune cell antigen staining over time in lung tissue from the High Dose MERS-CoV group. Altogether, these data suggest that increased MERS-CoV disease severity was associated with a potential failure in the adaptive immune response.

**Figure 5:**
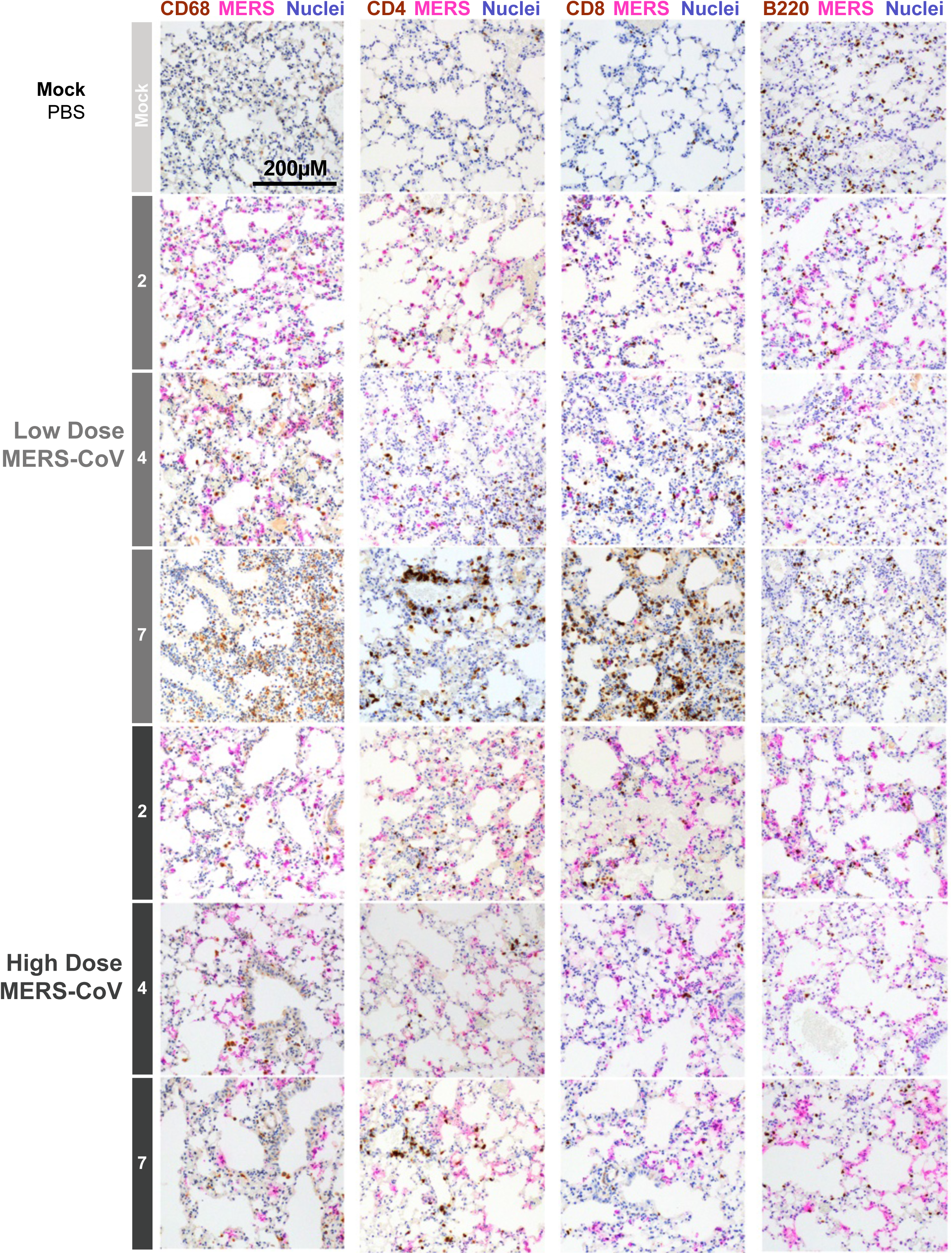
Severe MERS-CoV infection is associated with reduced innate and adaptive immune cells. Lung tissues from mock, low and high dose MERS-CoV infected mice were sectioned and immunostained for cellular nuclei (blue), MERS-CoV antigen (pink) and one of the following immune cell markers (brown): CD68 (monocyte/macrophage), CD4 (T cell subpopulation marker), CD8 (T cell subpopulation marker) or B220 (B-cell marker).

### Adaptive immune cells contribute to disease but are also important for controlling MERS-CoV replication

The collective data noted above demonstrated that adaptive immune signatures were dramatically different among those that do or do not survive MERS-CoV infection. To conclusively determine the importance of key adaptive immune cell populations in MERS-CoV pathogenesis, we generated a transgenic mouse susceptible to MERS-CoV infection lacking functional B and T cells through the crossing of C57BL/6 RAG^−/−^ and our C57BL/6 h288/330^+^DPP4 mice. C57BL/6 h288/330^+^DPP4 (WT) or C57BL/6 h288/330^+^DPP4 RAG^−/−^ (RAG^−/−^) mice were infected with Low or High dose MERS-CoV as described above after which body weight was monitored every day until 7 dpi when mice were humanely sacrificed, gross lung pathology was scored and lung tissue was harvested for viral lung titers (Fig. 6A). Regardless of virus dose, RAG^−/−^ mice lost significantly less weight than WT mice indicating that functional B or T-cells contribute to body weight loss (Fig. 6B). Although none of the Low Dose infected mice succumbed to infection, in the High Dose group, most of the WT (60%) and minority of RAG^−/−^ mice (30%) met human endpoints for euthanasia (Fig. 6C), suggesting that functional T or B cells contribute to severe disease. Remarkably, viral lung titers were significantly elevated in Low Dose MERS-CoV infected RAG^−/−^ mice over WT mice but were similar in both lines of mice in the High Dose group indicating that functional T and B cells were important in controlling MERS-CoV replication, but this phenotype was virus dose dependent (Fig. 6D). Congruent with the above data, gross lung pathology was elevated in High Dose MERS-CoV infected WT over RAG^−/−^ mice 7dpi (Fig. 6E). Similar trends were observed in follow up studies with a more pathogenic mouse adapted MERS-CoV strain m35C4 (Fig. 6S) (41). Collectively, these data demonstrate a multifunctional role for adaptive immune cells in MERS-CoV pathogenesis where functional T or B cells contribute to pathogenesis but also are important for controlling MERS-CoV replication.

**Figure 6:**
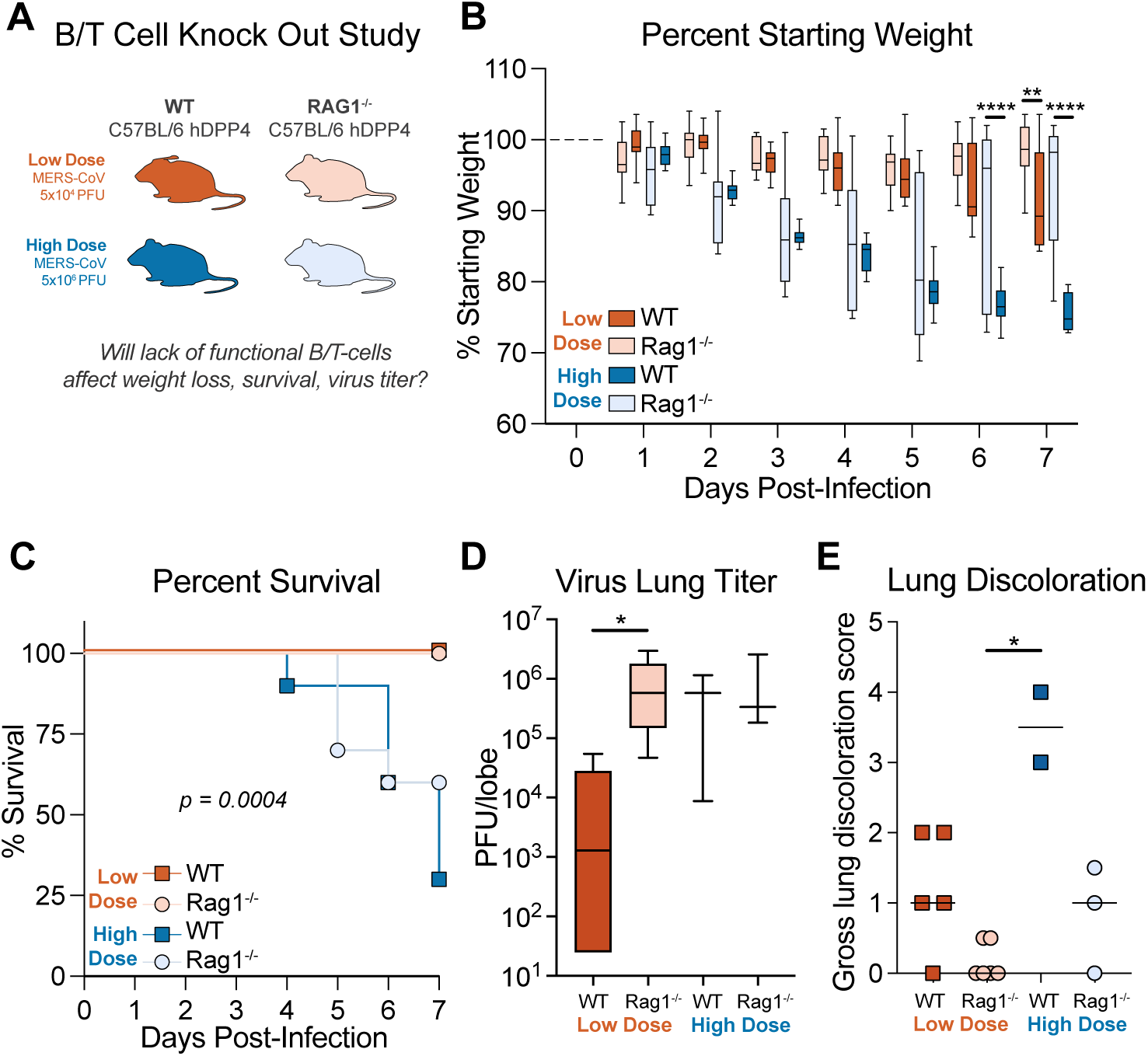
Adaptive immunity is protective and pathogenic in MERS-CoV disease. **(A)** Study design or MERS-CoV infection of mice deficient in functional T and B-cells. **(B)** Percent starting weight of 20-week old male and female WT C57BL/6 hDPP4 mice or C57BL/6 hDPP4 RAG1^−/−^ mice infected with 5×10^4^ PFU MERS ma1 (WT, N = 11; RAG1^−/−^ N = 10) or 5×10^6^ PFU MERS ma1 WT, N = 10; RAG1^−/−^ N = 12). Asterisks denote statistical significance as determined by Two-way ANOVA with a Tukey’s multiple comparison test. **(C)** Percent Survival. **(D)** Virus lung titer on 7dpi by plaque assay. Asterisks denote statistical significance as determined by Kruskal-Wallis test with a Dunn’s multiple comparison test. **(E)** Gross pathology “lung discoloration score” on 7dpi.

## Discussion

To date, there have been over 2,600 laboratory confirmed MERS-CoV cases and 943 deaths in 27 countries although the vast majority of cases have been in the Kingdom of Saudi Arabia (4). Serosurveys and epidemiologic investigations suggest that MERS-CoV infection is more widespread and may cause mild or asymptomatic disease especially in those younger than typical laboratory confirmed cases, and that those with mild or asymptomatic disease can be infectious and transmit virus (5, 42, 43). With noted region-specific genetic diversity, less pathogenic strains may circulate at low levels in East Africa, Pakistan and elsewhere (44). Risk factors for severe disease include direct contact with camels, increased age, male sex, diabetes, hypertension, cancer or lung disease (45). Similar to SARS-CoV-2, survivors of MERS-CoV can experience post-acute sequelae with radiographic evidence of lung fibrosis months after the acute phase of disease (46). Even though much has been learned about MERS-CoV pathogenesis since its discovery, the molecular mechanisms driving dichotomous clinical outcomes, the progression to severe disease and post-acute sequelae remains unclear. In addition, the threat level of *Merbecoviruses* remains high as many enzootic isolates replicate in primary human epithelial cells (47, 48).

Mouse models of emerging CoV pathogenesis have provided many insights into the viral and host genetic factors driving severe lung pathogenesis. For MERS-CoV, two residues (A288 in exon 10 and T330 in exon 11) in the murine ortholog (mDPP4) of the human receptor inhibit infection in mice but multiple genetic strategies have been employed to circumvent this issue (49). Pascal et al. replaced a majority the murine gene with that of human (exons 2-26), Li et al. replaced the murine region spanning exon 10 through 11 with those from hDPP4 and Cockrell et al. humanized only residues 288 and 330 via CRISPR/Cas9 (22, 50, 51). Given that DPP4 is a key mediator chemokine/cytokine responses and T-cell activation aside from being the receptor for MERS-CoV entry, chimeras of human and mouse DPP4 could affect infection, replication, pathogenesis and the immune response (13). The potential for this is evidenced by the fact that WT MERS-CoV EMC elicits ∼20% weight loss in the Pascal et. al model, yet Li et. al and Cockrell et. al had to passage virus to elicit severe disease phenotypes like ARDS and mortality (22, 51, 52). Interestingly, the host transcriptional responses noted in this report are like those reported by Coleman et. al using WT MERS-CoV in the Pascal et. al model (52). Thus, even though the pathogenic potential of WT MERS-CoV may vary by transgenic mouse model, mouse adapted MERS-CoV and WT viruses in these respective models appear to provide similar transcriptional disease signatures.

Emerging CoVs like SARS-CoV, SARS-CoV-2 and MERS-CoV can cause severe respiratory infection which can lead to ARDS. Cellular entry receptor and expression in the respiratory tract likely plays a role in transmissibility, pathogenesis, disease severity and recovery. Unlike SARS-CoV-2 which primarily targets AT2 cells in the terminal airway, MERS-CoV predominantly targets both AT1 and AT2 cells and replicates efficiently in primary human lung fibroblasts and endothelial cells (16, 20, 45). Over 95% of the alveolar surface area is made up of the morphologically flat AT1 cells which form a physical barrier, structure the alveoli and mediate gas exchange through alveolar capillaries (53). In contrast, AT2 cells are cuboidal, are less abundant in the terminal airways but are the main producers of surfactant, the lipoprotein mixture that reduces alveolar surface tension without which lungs would collapse (54). Here, we show that **i**) MERS-CoV infection of AT1 and AT2 cells results in the induction of apoptosis; **ii)** severe disease is associated with lung pathology consistent with ARDS; **iii)** severe disease is associated with a reduction in AT1 and AT2 cell associated gene and protein expression and lipid production; and **iv)** a reduction in surfactant protein expression that does not return to baseline by 7dpi with severe MERS-CoV disease. Altogether, these data demonstrate that MERS-CoV infection has a profound effect on function and viability of AT1 and AT2 cells, which together play essential roles in maintaining homeostasis and pulmonary function. In mouse models of acute and post-acute sequelae of COVID-19 (PASC), a reduction in surfactant expressing cells was also observed early and was associated with the development of lung fibrosis in those that survive severe acute infection (55). Survival of severe SARS-CoV-2 infection in mouse and human can lead to the initiation of dysregulated repair programs and chronic pneumonias with fibrotic lesions and other abnormalities. Although speculative, there is evidence in humans to support a similar hypothesis for other emerging CoV. Thirty five percent of MERS-CoV survivors have abnormal chest radiographs characterized by lung fibrosis (33%), ground glass opacities (5.5%) and pleural thickening (5.5%) (46). In a small longitudinal cohort of SARS-CoV survivors in Hong Kong, diminished pulmonary function was observed in almost 25% of patients a year after illness onset (56). Similarly, in a cohort of survivors of the 2015 outbreak in South Korea, those that experienced more impaired pulmonary function during acute disease had more impaired pulmonary function after a year which was confirmed radiologically (57). Interestingly, we and others recently demonstrated that antiviral therapy during the acute phase of SARS-CoV-2 infection diminished PASC phenotypes in mice and that pharmacologic inhibition of PERK-mediated apoptosis in MERS-CoV infected mice improved short term outcomes (21, 55, 58). Therefore, direct acting antivirals as well as modulators of the host response could be used to improve outcomes of both acute and post-acute MERS-CoV disease outcomes, a subject of future investigation.

Studies of immune responses in MERS-CoV patients in Saudi Arabia and South Korea demonstrate that high viral loads, lymphopenia and thrombocytopenia as well as high neutralizing antibody titers and higher numbers of virus specific CD4 T-cells were associated with increased disease severity and death implicating a role for both humoral and cell-mediated immunity in MERS-CoV disease (59, 60). Interestingly, as DPP4 is expressed on immune cells aside from the main epithelial cell targets, MERS-CoV infection of human T-cells ex-vivo has been reported leading to the induction of cell death pathways (61). MERS-CoV encodes antagonists of innate immunity and infection is associated with a downregulation of certain interferon stimulated genes and proteins that mediate antigen presentation thus demonstrating the importance of the interplay among innate and adaptive immunity in controlling MERS-CoV infection (62, 63). Like that observed in humans, here we show that innate and adaptive immune related genes are downregulated with severe MERS-CoV. Since we performed “bulk” omics analysis, we cannot determine if these signatures originated in infected or bystander cells. In the Pascal et al. model, Coleman et al. demonstrated that antibody depleting CD8+ but not CD4+ cells reduced MERS-CoV pathogenesis but neither treatment affected lung viral loads (52). Similarly, Channappanavar et. al performed similar studies in the Li et al. model simultaneously depleting both CD4+ and CD8+ cells with antibodies which resulted in suboptimal viral clearance and significant viral titers remaining at 7dpi (64). Rather than take an antibody depletion approach which is transient and can be incomplete, we leveraged mouse transgenics through the breeding of our MERS hDPP4 with RAG1^−/−^ mice, which lack functional T and B cells. Infection of the resultant double transgenic mice (C57BL/6 hDPP4 RAG1^−/−^) with two different mouse adapted MERS-CoV viruses resulted in diminished pathogenesis and significantly increased viral lung titers 7dpi implicating T and/or B cells were important for both the control of virus replication and driving severe disease. Collectively, data from multiple DPP4 transgenic mouse models support that adaptive immunity likely contributes to both protective and pathogenic responses to MERS-CoV infection yet the molecular mechanisms responsible for this dichotomy are not completely understood (22, 51).

As three novel CoVs have emerged in the past 20 years and many enzootic *Merbecoviruses* and *Sarbecoviruses* are poised for emergence, it is likely novel CoVs will continue to emerge causing novel human disease (47, 48, 65, 66). The comprehensive disease signatures generated through multi-omics studies of viral pathogenesis offer a unique opportunity to not only better understand the molecular mechanisms of disease but also to identify genes and pathways that can be exploited for therapeutic intervention all of which is important for our future pandemic preparedness.

## Materials and Methods

### Viruses and Cells

Infectious clone derived mouse adapted MERS-CoV MA1 (MERS-CoV MA1)(22) and mouse adapted plaque purified clone MERS-CoV m35C4(41) stocks were grown in Vero CCL81 cells (ATCC, CCL81). MERS-CoV MA1 differs from parental MERS-CoV EMC/2012 in 5’ UTR (ΔA, nucleotide 2), nsp3 A217V, nsp6 T184I, nsp8 I108L, spike R884-RMR insertion, spike S885L, ns4b from nt 26,226 to 26,821(ΔE45-H243) (22). MERS-CoV MA2 differs from parental MERS-CoV EMC/2012 in 5’ UTR (ΔA, nucleotide 2), nsp2 A196V, nsp6 T184I, nsp8 I108L, spike R884-RMR insertion, spike S885L, ns4b from nt 26,211 to 26,863(ΔS40 in ORF4b to 8F in ORF5) (22). MERS-CoV m35C4 is a highly pathogenic further mouse adapted virus with MERS-CoV MA1 as parent. MERS-CoV m35C4 is different than MERS-CoV EMC/2012 at: 5’ UTR (A:G, nucleotide 28), nsp3 A217V, nsp3 T1615N, nsp6 L232F, nsp8 I108L, nsp14 T521I, spike N222Y, spike R884-RMR insertion, spike S885L, orf3 Q4 stop codon, orf4a P85L, orf4b/orf5 deletion (nucleotide 26211-26863), orf M S2F(41). CCL81 cells were maintained routinely in Dulbecco’s modified Eagle’s medium (Gibco) supplemented with 10% fetal bovine serum (FBS; Hyclone) and 1x antibiotic/antimycotic (Gibco). All viruses were harvested in Opti-MEM medium (Gibco) supplemented with 3% FBS, 1x antibiotic/antimycotic (Gibco), 1x non-essential amino acids (Gibco) and 1x 1mM sodium pyruvate (Gibco).

### Biosafety

All work for these studies was performed with approved standard operating procedures and safety conditions for MERS-CoV (not a select agent). Our institutional CoV BSL3 facilities have been designed to conform to the safety requirements that are recommended by the Biosafety in Microbiological and Biomedical Laboratories, the US Department of Health and Human Services, the Public Health Service, the Centers for Disease Control and the NIH. Laboratory safety plans were submitted to, and the facility has been approved for use by, the UNC Department of Environmental Health and Safety (EHS) and the CDC. All workers have been trained by EHS to safely use powered air-purifying respirators, and appropriate work habits in a BSL3. Our BSL3 facilities contain redundant fans, emergency power to fans, and biological safety cabinets and freezers, and our facilities use SealSafe mouse racks. Within the BSL3 facilities, experimentation with infectious virus is performed in a certified Class II Biosafety Cabinet, linked to emergency power. All members of staff wear scrubs, Tyvek suits and aprons, powered air-purifying respirators and shoe covers, and their hands are double-gloved. BSL3 users are subject to a medical surveillance plan monitored by the University Employee Occupational Health Clinic (UEOHC), which includes a yearly physical, annual influenza vaccination and mandatory reporting of any symptoms associated with coronavirus infection during periods when working in the BSL3. All BSL3 users are trained in exposure management and reporting protocols, are prepared to self-quarantine and have been trained for safe delivery to a local infectious disease management department in an emergency. All potential exposure events are reported and investigated by EHS and UEOHC, with reports filed to both the CDC and the NIH.

### Ethics statement

Mouse studies were carried out in accordance with the recommendations for the care and use of animals by the Office of Laboratory Animal Welfare at NIH. IACUC at UNC-CH approved the animal studies performed here (protocol, IACUC 16-251), using a weight loss cut-off point of ∼30% for humane euthanasia.

### MERS-CoV infection and pathogenesis in mice for multi-omics analysis

Ketamine and xylazine anesthetized, 19-23 week old female C57BL/6 288/330+/+ mice (C57BL/6 hDPP4) were intranasally infected with PBS (“mock”, N = 14), 5×10^4^ PFU mouse adapted MERS ma1 (“low dose”, N = 14) or 5×10^6^ PFU MERS ma1 (“high dose”, N = 16) in a 50µl volume. Body weight loss, a crude marker of emerging coronavirus pathogenesis, was recorded daily. Virus induced mortality was recorded for animals found dead in cage or if animals met criteria for humane euthanasia by isoflurane overdose (30% loss in starting weight, see below). On 2, 4, and 7dpi, 4 mice per group were humanely sacrificed by isoflurane overdose. Lung hemorrhage was first scored on a scale of 0–4, where 0 is a normal pink healthy lung and 4 is a completely dark red lung(26). Lung tissue was then harvested for virus titer (left superior lobe), pathological (right lobe), transcriptomic (left middle and post caval lobes) and proteomic, lipidomic and metabolomics (inferior left lobe) analysis to be discussed in detail below. Virus titer was determined by plaque assay in Vero CCL81 cells as described(26). Briefly, medium was removed from 6-well plates seeded with 500,000 Vero CCL81 the day prior, and serial dilutions of sample were added per plate (10−1–10−6 dilutions) and incubated at 37 °C for 1 h, after which wells were overlayed with 1× DMEM, 5% Fetal Clone 2 serum, 1× A/A, 0.8% agarose. Three days after, plaques were enumerated to generate a plaque/ml value.

### Transcriptomics, proteomics and lipidomics sample processing

Immediately after removal from the mouse, the left middle and post caval lobes of lung tissue for transcriptomics was stored in RNAlater (Invitrogen, AM7021) at −80°C until processing by homogenization in Trizol (Invitrogen/ThermoFisher) followed by isolation of total RNA according to the Trizol protocol. Total RNA samples were sent to Arraystar for microarray analysis. In parallel, the inferior left lobes were subjected to the MPLEx protocol (chloroform/methanol, 2:1 mix) precipitation as previously described(67–69), which results in simultaneous extraction of proteins, metabolites, and lipids with concomitant inactivation of virus. All proteomic and lipidomic samples were desiccated and frozen prior to shipment to Pacific Northwest National Laboratories for downstream analysis.

### Transcriptomic Analysis

Scanned images were analyzed using Agilent Feature Extraction Software (v11.0.1.1). The limma package for R (available on Bioconductor) was used to perform background correction, quantile normalization (normalizeBetweenArrays), and summarization (avereps) to derive a single normalized intensity value per probe. Outlier samples were detected using PCA and by visual inspection of heatmaps, and all data was re-processed after removing outlier samples. All data processing for each of the biological replicates was performed independently of the other.

### Proteomic Analysis

Samples for proteomics analysis were prepared for and analyzed using liquid chromatography-mass spectrometry and the Accurate Mass and Time Tag approach, as described previously (70). The RMD-PAV algorithm (71) was used to identify any outlier biological samples and was confirmed via Pearson correlation. Peptides with inadequate data for either qualitative or quantitative statistical tests were also removed from the dataset(72). The SPANS algorithm (72) was used to identify the best normalization method for each dataset. Peptides were evaluated with Analysis of Variance (ANOVA) with a Dunnett test correction and a Bonferroni-corrected g-test to compare each virus to the associated mock within each time point. To perform signature-based protein quantification, BP-Quant(73), each peptide was categorized as a vector of length equal to the number of viruses being evaluated. If all comparisons for all time points are 0 for a specific virus it is considered as non-changing and given a value of 0. If there are more time points with an increase in virus to mock than decreasing it is categorized as a +1 and the contrary −1 is given for the decrease in virus to mock. BP-Quant was run with a default parameter of 0.9. All proteins were then analyzed using the same methodology as for the peptides; ANOVA with a Dunnett test correction and a Bonferroni-corrected g-test to compare each virus to the associated mock within each time point.

### Lipidomic Analysis

Samples for lipidomics analysis were prepared for and analyzed using liquid chromatography-tandem mass spectrometry, as described previously (55). Lipids were identified using the in-house tool LIQUID (86), and their quantitative data were extracted using MZmine 2.0 as previously described (87). The RMD-PAV algorithm (83) was also used to identify any outlier biological samples and was confirmed via Pearson correlation. Lipids with inadequate data for either qualitative or quantitative statistical tests were also removed from the dataset (84). Median centering was used for normalization. Lipids were evaluated with a standard two sample t-test to compare each infected condition to the associated mock within each time point.

### MERS-CoV pathogenesis studies in RAG1^−/−^ mice

To determine the importance of functional T-cells and B-cells on MERS-CoV pathogenesis, we bred C57BL/6 hDPP4 mice with C57BL/6 RAG1^−/−^ (Jackson Labs Strain #002216) mice resulting in C57BL/6 hDPP4 RAG1^−/−^ offspring. 20-week old male and female WT C57BL/6 hDPP4 mice or C57BL/6 hDPP4 RAG1^−/−^ mice infected with 5×10^4^ PFU MERS ma1 (WT, N = 11; RAG1^−/−^ N = 10) or 5×10^6^ PFU MERS ma1 (WT, N = 10; RAG1^−/−^ N = 12) after which weight loss, mortality and virus titer was monitored over time. We then evaluated the pathogenic potential of the further adapted MERS-CoV m35c4 virus in WT C57BL/6 hDPP4 mice or C57BL/6 hDPP4 RAG1^−/−^ mice (WT, N = 11; RAG1^−/−^ N = 10) after which weight loss, mortality and virus titer was monitored over time.

### Lung pathology and acute lung injury scoring

Formalin fixed and paraffin embedded lung tissue sections (5µm) were stained with hematoxylin and eosin and blindly evaluated by a board-certified veterinary pathologist. Two complementary tools were employed to quantitate the histological features of acute lung injury. First, we utilized a tool created by the American Thoracic Society (ATS)(28). In three random diseased fields of lung tissue at high power (60×), the following was scored: (A) neutrophils in the alveolar space (none, 0; one to five cells, 1; >5 cells, 2), (B) neutrophils in the interstitial space/septae (none, 0; one to five cells, 1; >5 cells, 2), (C) hyaline membranes (none, 0; one membrane, 1; >1 membrane, 2), (D) proteinaceous debris in air spaces (none, 0; one instance, 1; >1 instance, 2), and (E) alveolar septal thickening (<2× mock thickness, 0; 2 to 4× mock thickness, 1; >4× mock thickness, 2). To obtain a lung injury score per field, the scores for A to E were then put into the following formula, which contains multipliers that assign varying levels of importance for each phenotype of the disease state: score = [(20 × A) + (14 × B) + (7 × C) + (7 × D) + (2 × E)]/100. The scores for the three fields per mouse were averaged to obtain a final score ranging from 0 to and including 1. The second tool specifically examines diffuse alveolar damage (DAD), the pathological hallmark of ALI(28, 74). Three random diseased fields of lung tissue were scored at high power (60x) for the following in a blinded manner: 1, absence of cellular sloughing and necrosis; 2, uncommon solitary cell sloughing and necrosis (one to two foci per field); 3, multifocal (3 + foci) cellular sloughing and necrosis with uncommon septal wall hyalinization; or 4, multifocal (>75% of field) cellular sloughing and necrosis with common and/or prominent hyaline membranes. The scores for the three fields per mouse were averaged to get a final DAD score per mouse.

### RNA *in situ* hybridization/Quantification

RNA-ISH was performed on paraffin-embedded 5 μm tissue sections using the RNAscope Multiplex Fluorescent Assay v2 according to the manufacturer’s instructions (Advanced Cell Diagnostics). Briefly, tissue sections were deparaffinized with xylene and 100% ethanol twice for 5 min and 1 min, respectively, incubated with hydrogen peroxide for 10 min and in boiling water containing target retrieval reagent for 15 min, and then incubated with Protease Plus (Advanced Cell Diagnostics) for 15 min at 40°C. Slides were hybridized with custom probes at 40°C for 2 h, and signals were amplified according to the manufacturer’s instructions. Stained sections were scanned and digitized by using an Olympus VS200 fluorescent microscope. Images were imported into Visiopharm Software^®^ (version 2020.09.0.8195) for quantification. Lung tissue and probe signals for targeted genes were quantified using a customized analysis protocol package to 1) detect lung tissue using a decision forest classifier, 2) detect the probe signal based on the intensity of the signal in the channel corresponding to the relevant probe. All slides were analyzed under the same conditions. Results were expressed as the area of the probe relative to total lung tissue area.

|  |  |  |
| --- | --- | --- |
| RNAScope 2.5 HD Duplex Reagent Kit | ACD | Cat#322430 |
| RNAScope Multiplex Fluorescent Reagent Kit v2 | ACD | Cat#323100 |
| RNAScope probe V-MERS-CoV-O1 (channel 1) | ACD | Cat#485081 |
| RNAScope probe V-MERS-CoV-S (channel 3) | ACD | Cat#860551-C3 |
| RNAScope probe Ager (channel 1) | ACD | Cat#550791 |
| RNAScope probe Sftpc (channel 2) | ACD | Cat#314101-C2 |
| RNAScope probe Foxj1 (channel 2) | ACD | Cat#317091-C2 |
| RNAScope probe Scgb1a1 (channel 3) | ACD | Cat#420351-C3 |

### Immunohistochemistry

Immunohistochemical staining was performed on paraffin-embedded 5 μm tissue sections according to a protocol as previously described (75). Briefly, paraffin-embedded sections were baked at 60 °C for 2– 4 hours, and deparaffinized with xylene (2 changes × 5 min) and graded ethanol (100% 2 × 5 min, 95% 1 × 5 min, 70% 1 × 5 min). After rehydration, antigen retrieval was performed by boiling the slides in 0.1 M sodium citrate pH 6.0 (3 cycles with microwave settings: 100% power for 6.5 min, 60% for 6 min, and 60% for 6 min, refilling the Coplin jars with distilled water after each cycle). After cooling and rinsing with distilled water, slides were blocked with Blocking One Histo (Nacalai Tesque) for 20 min at RT. Primary antibodies (MERS anti sera: 1:500, Ager: 1:400, Lamp3: 1:100, Cleaved caspase-3: 1:200) were diluted in Blocking One Histo (1:20 in PBST) and applied to the slides, incubated over night at 4 °C. Mouse, goat, rat, and rabbit gamma globulin were used as an isotype control at the same concentrations as the primary antibodies. Sections were washed in PBST and species-specific secondary antibodies were applied for 60 min at RT. After washing in PBST, the Vector® TrueVIEW Autofluorescence Quenching Kit (Vector Laboratories) was used to reduce non-specific background staining, and glass coverslips were placed over tissue sections with the Vectashield Vibrance with DAPI (Vector Laboratories). Coverslipped slides were scanned and digitized using an Olympus VS200 fluorescent microscope.

|  |  |  |
| --- | --- | --- |
| Goat polyclonal Ager | R&D Systems | RRID: AB_354628 |
| Rabbit Cleaved caspase-3 | Cell Signaling | RRID: AB_2341188 |
| Rat DC-Lamp (Lamp3) | Novus Biologicals | RRID: AB_2827532 |
| Mouse MERS anti sera |  |  |

### Multi-Omics data availability

The multi-omics datasets described here in detail were generated under the ‘Omics of Lethal Human Viruses (OMICS-LHV) Systems Biology Center was funded by the National Institutes of Allergy and Infectious Diseases (NIAID) (76). The accession numbers for the above data sets are publicly available here: Transcriptomics: GSE108594 (GEO) Proteomics: MSV000083532 (MassIVE) Lipidomics: MSV000083536 (MassIVE)

## Acknowledgements

This study was funded by NIH/NIAID U19 AI109761, and NIH U19 AI106772 grants. A portion of the research described in this paper was conducted under the Laboratory Directed Research and Development Program at Pacific Northwest National Laboratory, a multiprogram national laboratory operated by Battelle for the U.S. Department of Energy. A portion of the research was performed using the Environmental Molecular Sciences Laboratory, a national scientific user facility sponsored by the U.S. Department of Energy’s Office of Biological and Environmental Research and located at Pacific Northwest National Laboratory (PNNL). PNNL is operated by Battelle Memorial Institute for the U.S. Department of Energy under Contract DE-AC05-76RL01830.

## Author contributions

ACS, ASC, TPS, RSB, TOM, KMW designed in vivo studies to obtain multi-omics samples. TPS, ACS, ASC, JEK, KEBJ, TOM, and KMW harvested, isolated and/or processed systems biology samples and/or datasets. SRL, AS, JFK, ASC, KLJ and TPS designed and executed knock out mouse studies. SAM evaluated all histology, lung injury scoring and immune cell immunohistochemistry. KO performed and analyzed all in situ hybridization and epithelial cell population immunohistochemistry. TPS, RSB, ACS, RCB, KO and SRL wrote and edited the manuscript. RSB, KMW, RCB, RDS, TOM and KMW raised funds for these studies.

## Supplemental Figure Legends

**Figure 1S: Levels of MERS-CoV viral antigen is dose dependent in lung tissue sections**. Related to Main Figure 1. Lung tissue sections from the study described in Main Figure 1 were labeled for MERS-CoV antigen (brown) and counterstained with hematoxylin (blue). Representative images from 2, 4 and 7 dpi are shown.

**Figure 2S: Surfactant expression is diminished with severe MERS-CoV disease and is not recovered by 7dpi**. RNAscope in situ hybridization for MERS-CoV and *Sftpc* RNA for mock, Low and High dose infection conditions at 2, 4 and 7dpi. **(A)** Quantitation of *Sftpc* RNA labeled using RNAscope. RNAscope in situ hybridization for MERS-CoV and *Sftpc* RNA for mock (**B)**, Low and High dose infection conditions at 2, 4 and 7dpi **(C).**

**Figure 3S: MERS-CoV disease severity is associated with modulation of alveolar epithelial cell transcriptomic, proteomic and lipidomic signatures.** Transcriptomic **(A)** and proteomic **(B)** data was filtered through epithelial cell signatures identified through single cell RNAseq by Treutlein et al. (33). Heat maps show log_2_ fold-change over mock data and the color bar indicates the epithelial cell type associated with the pattern. **(C)** MERS-CoV disease severity is associated with increased modulation of the global lipidome. Heatmap shows the log_2_ fold-change data for significantly regulated lipids in lung tissue. Significance was determined by ANOVA (P = 0.05) when enough data was present or g-test for qualitative comparisons and the times at which lipids are significantly regulated are plotted in the heatmap on the right. **(D)** Surfactant and inflammation associated lipids. Log_2_ fold-change data for CE, CER, LPC, PC, LPI, PG and TG. Shorter-chain PC and PG are main surfactant lipids and shorter-chain TG act as a fatty acid reservoir for surfactant producing cells. LPC, LPI and long chain TGs are associated with inflammation.

**Figure 4S: Most innate immune related gene expression is similar across MERS-CoV dose groups. (A-B)** Significantly regulated transcripts and proteins for innate immune response related canonical pathways from **“Figure 4”**: **(A)** Interferon Signaling and **(B)** Recognition of Viruses by Pattern Recognition Receptors **(B)**. **(C-D)** Transcriptomic and proteomic data was filtered through a list of 390 known interferon stimulated genes (ISGs) (77). **(C)** Heat map of the differentially regulated ISG transcripts **(C)** or proteins **(D)** (log_2_ fold-change) for both high and low dose MERS-CoV infection. Gene ontology information (biological function, positive/negative regulatory role) is also noted.

**Figure 5S: High dose MERS-CoV infection is associated with a downregulation of adaptive immune gene and protein expression.** Significantly regulated transcripts and proteins for adaptive immune response related canonical pathways from **“Figure 4”:** Th1 Pathway **(A)**, NFAT Regulation of Immune Response **(B)**, Leukocyte Extravasation Signaling **(C)**, and p70S6K Signaling **(D)**. All heat maps show log_2_ fold-change over mock data per gene/protein significantly affected for each pathway.

**Figure 6S: Functional T and/or B-cells contribute to MERS-CoV disease and are important for controlling MERS-CoV replication**. Related to Main Figure 6. **(A)** Percent starting weight of 10-week old male and female WT C57BL/6 hDPP4 mice or C57BL/6 hDPP4 RAG1^−/−^ mice infected with 2.4×10^4^ PFU of the further mouse adapted MERS m35C4 strain (WT, N = 11; RAG1^−/−^ N = 10). Asterisks denote statistical significance by two-way ANOVA with a Sidak’s multiple comparison test. **(B)** Percent Survival. **(C)** Gross lung pathology “lung discoloration”. Asterisks denote statistical significance as determined by Wilcoxon matched-pairs signed rank test. **(D)** Virus lung titer on 7dpi by plaque assay. Asterisks denote statistical significance as determined by Mann Whitney test.

